# Phenylephrine induces relaxation of longitudinal strips from small arteries of goat legs

**DOI:** 10.1101/2019.12.18.881003

**Authors:** Kawin Padmaja Marconi, Bhavithra Megan, Alen Major Venis, Renu Raj, Sathya Subramani

## Abstract

Alpha adrenergic stimulation is known to produce vasoconstriction. We have earlier shown that, in spiral strips of small arteries Phenylephrine (PE) caused vasorelaxation under high nitric oxide (NO) environment. However on further experimentation it was realized that the PE-induced vasorelaxant response occurred only with longitudinal strips of small arteries even under normal NO environment while circular strips showed contraction with PE even under high NO environment. Such PE-induced vasorelaxation of longitudinal strips was blocked by Phentolamine, an alpha-adrenergic receptor blocker. On delineation of specific receptor subtype, PE-induced relaxation was found to be mediated through alpha 1D receptor. However, this phenomenon is specific to small artery, as longitudinal smooth muscle of aorta showed only contractile response to adrenergic stimulation. There is no prior report of longitudinal smooth muscle in small artery up to our knowledge. The results of this study and histological examination of vessel sections suggest the presence of longitudinal smooth muscle in small artery and their relaxant response to alpha adrenergic stimulation is a novel phenomenon.

## INTRODUCTION

Phenylephrine is an alpha-adrenergic agonist which is known to cause contraction of vascular smooth muscle. We had recently reported that Phenylephrine caused relaxation of spiral strips made from small arteries from goat legs in a high nitric oxide environment, while causing contraction in normal circumstances. However, with further experimentation we have realized that longitudinal strips made from small arteries always relaxed with Phenylephrine, even in normal nitric oxide environment. On the other hand, PE caused contraction of circular strips of small arteries even in a high nitric oxide environment. Spiral strips used in earlier experiments have led to erroneous inferences, as the method of making the strip is prone to experimenter bias.

In this paper, we compare the responses of transverse (or circular) and longitudinal preparations made from aorta and small arteries from goats. While PE increases vascular tension in both transverse and longitudinal strips made from aorta, as do circular strips of small arteries, longitudinal strips from small arteries show reduction in vascular tension in the presence of PE.

An important determinant of arterial pressure is peripheral vascular resistance, which in turn is primarily determined by the diameter of small arteries and arterioles which are referred to as resistance vessels (1). The walls of the resistance vessels have smooth muscle cells arranged in a concentric fashion in the tunica media which is referred to as circular smooth muscle (2). Contraction of circular smooth muscle reduces lumen diameter and therefore increases vascular resistance (1) (3). The balance between vasoconstrictor and vasodilator signals acting on the circular smooth muscle determines the diameter and therefore the tone of resistance vessels.

While muscular arteries (medium sized) are documented to have concentrically arranged circular smooth muscle in the tunica media, veins have longitudinal smooth muscle just next to the intima (4). There is evidence for the presence of longitudinal smooth muscle in the walls of large arteries and aorta too(5). Function of longitudinal smooth muscle in the large arteries has not been clearly understood.

There is, however, no prior report of longitudinal smooth muscle in small arteries, up to our knowledge. The results of this study suggest the presence of longitudinal smooth muscle in small arteries, and their relaxant response to alpha adrenergic stimulation is a novel phenomenon.

## Materials & Methods

### Solutions required for the experiment

The composition of the mammalian ECF solution used in the experiments was as follows (in mmol/L): NaCl 100; KCl 3; CaCl_2_ 1.3; NaH_2_PO_4_ 0.5; Na_2_HPO_4_ 2; NaHCO_3_ 25; MgCl_2_ 2; HEPES 10; Glucose 5; pH was corrected to 7.4 with 1mol/L sodium hydroxide. All salts for mammalian ECF solution were purchased from SIGMA. Phenylephrine hydrochloride, L-Arginine, Sodium Nitroprusside (SNP) and L-NNA were also purchased from SIGMA.

10 mmol/L stock solution was prepared for phenylephrine and L-Arginine in distilled water. Appropriate amount of the drug was added both to the organ bath and to the drug reservoir to obtain the final concentration.

### Isolation of small sized artery

Fresh goat leg from a registered slaughterhouse (latitude and longitude) was procured on the day of experiment, washed, and the skin removed. A vascular bundle close to the muscle was identified. The artery in the bundle was identified by patency of its lumen. Considerable length of the artery was dissected and transferred to a petri dish containing cold ECF solution. Adventitious tissue attached to the artery was removed and the artery was cut into segments of 1.5 – 2 cm length. It was ensured that the arterial segments were devoid of side branches. Two different arterial preparations were made:

i. Transverse cylinder
ii. Longitudinal strip

### Transverse cylinder preparation

Two fine threads were inserted through the lumen of the arterial segment and loops were made on opposite sides. One loop was connected to a force transducer and the other loop was attached to a hook in the organ bath such that the tissue was suspended transversely. (Fig.1)

**Fig.1:**
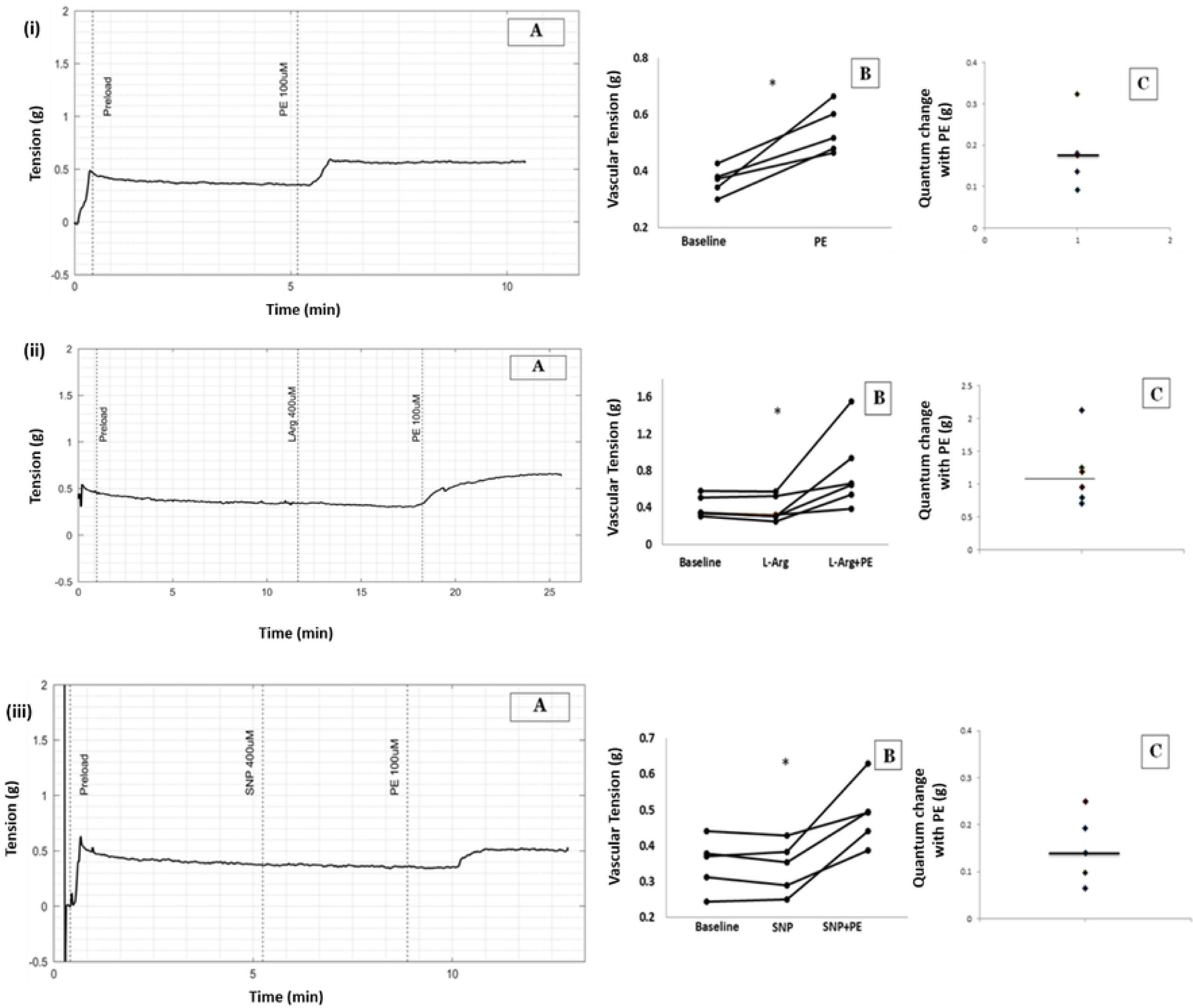
Diagrammatic representation of transverse cylinder preparation and longitudinal strip preparation of small artery

### Longitudinal strip preparation

An arterial segment of length 1.5 – 2 cm was selected and cut open longitudinally using an iris scissors thereby exposing the endothelium. One end of the longitudinal axis was connected to the force transducer while the other end was connected to the base of the organ bath. (Fig.1)

### Isolation of aorta

Fresh goat heart procured from a registered slaughterhouse was transported to lab in ice cold mammalian ECF solution. The heart was washed, pericardial sac and fat removed, and ascending aorta was identified by the patency of its lumen. The aorta was then dissected from its point of origin to the point before it branches. The dissected aorta was transferred to a petri dish containing cold ECF solution and the adventitious tissue removed. Two different preparations were made.

i. Aortic circular strip
ii. Aortic longitudinal strip

### Aortic circular strip preparation

A ring of aorta was cut, with thickness of 5 mm. One end of the ring was attached to the force transducer and the other end to the base of the organ bath. (Fig.2)

**Fig.2:**
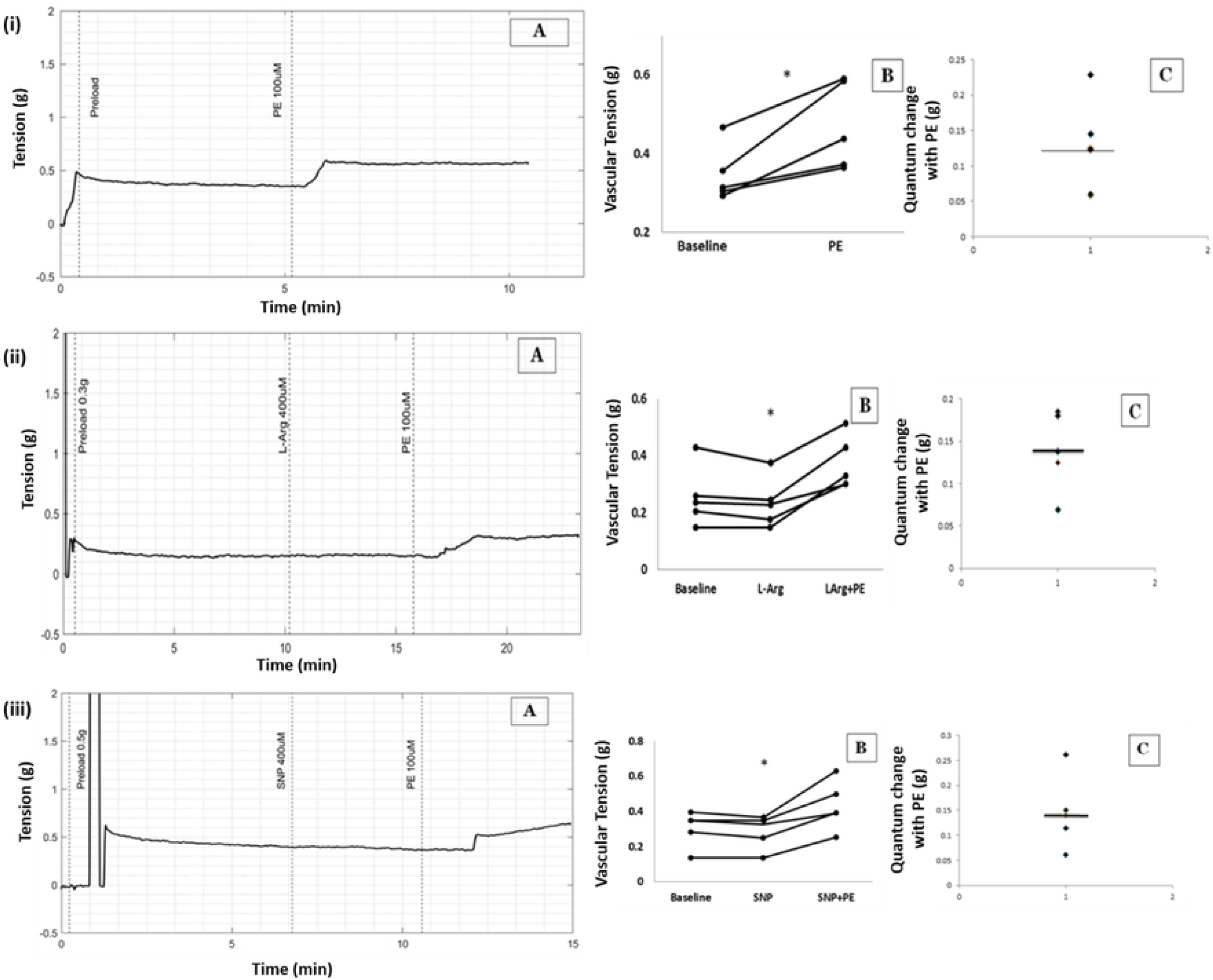
(A) Raw tracings showing changes in vascular tension in circular strips of aorta (B) Scatter plots of results from all five experiments demonstrating increase in vascular tension with PE. (C) Quantum of change with PE from baseline (* p< 0.05). (i) PE 100uM (ii) PE 100uM in high NO environment caused by L-Arginine (iii) PE 100uM in presence of SNP

### Aortic Longitudinal Preparation

A length of aortic cylinder was cut open in longitudinal orientation with surgical scissors exposing the endothelium. Opposite ends of the longitudinal strip were connected to a force transducer and the base of the organ bath.

The force transducer was connected to Powerlab data acquisition system for recording vessel tension. The thread was kept taut with an optimum preload and the resting tension was maintained between 0.2 to 0.6g. Change in tension was recorded when drugs were added into the organ bath. The data was acquired at 1 KHz and was processed using MATLAB R2018a software.

### Histological examination of vessel preparations

Transverse and longitudinal preparations of small artery and aorta were immersed in 10% buffered formalin for 2-3 days for fixation. Ascending grades of alcohol were used to dehydrate the tissues which were finally cleared with xylene. Then paraffin blocks of the tissues were made by impregnating with liquid paraffin. 5µm thickness sections were made using microtome. Tissue sections were subjected to dewaxination and were hydrated with descending grades of alcohol and were washed in distilled water. Staining was done using hematoxylin and eosin and sections were observed under light microscope.

### Statistical analysis

Statistical analysis was done using SPSS software ver.16.0. Comparison of vascular tension before and after intervention in same group was done using Wilcoxon signed rank (WSR) test. Changes in vascular tension between different groups were done using Mann-Whitney U test. P value ≤ 0.05 was considered statistically significant.

## RESULTS

A summary of the major results obtained is presented in the table below and the details are provided subsequently:

|  | **Aorta**<br>**circular strip** | **Aorta**<br>**longitudinal**<br>**strip** | **Small artery**<br>**transverse**<br>**Strip** | **Small artery**<br>**longitudinal**<br>**strip** |
| --- | --- | --- | --- | --- |
| **PE 100 μM** | Contraction | Contraction | Contraction | Relaxation |
| **L-Arginine + PE** | Contraction | Contraction | Contraction | Relaxation |
| **SNP+ PE** | Contraction | Contraction | - | - |
| **L-NNA + PE** | - | - | No Contraction | No Relaxation |
| **Phentolamine + PE** | - | - | No Contraction | No Relaxation |

### PE induces contraction of circular smooth muscles of aorta under Normal NO and high NO environments as expected

In circular strips of aorta, PE 100 µM increased tension from 0.37 to 0.51g. (Median, n = 5, P value = 0.043 with WSR test comparing pre and post PE tension), Fig: 2, (i).

L-Arginine was added to the bath to increase NO levels in the circular strip of aorta. No change in tension was observed with L-Arginine. Subsequent addition of PE 100 µM increased tension from 0.31g to 0.65g. (Median, n=6, P value = 0.028 with WSR test, Fig.2, (ii).

Addition of sodium nitroprusside (SNP) 400uM to the bath to create high NO environment, did not change vascular tension in the circular aortic strip. Subsequent addition of PE increased tension from 0.35g to 0.49g. (Median, n=5, P value = 0.043 with WSR when tension before and after in presence of SNP were compared), Fig.2, (iii).

There was no significant difference when percentage change in aortic vascular tension due to PE with and without L-Arginine (P value = 0.273 with MWU) were compared and also when PE with and without SNP (P value = 0.754 with MWU) were compared, showing that PE-induced vasoconstriction in aortic circular muscle was not affected by high levels of NO.

### PE induces contraction of longitudinal smooth muscle of aorta under normal and high NO environment

PE 100uM increased the tension of longitudinal smooth muscle in aorta from 0.31g to 0.43g. The increase in tension by PE was statistically significant. (Median, n=5, P value <0.05, WSR), Fig.3, (i).

**Fig.3:**
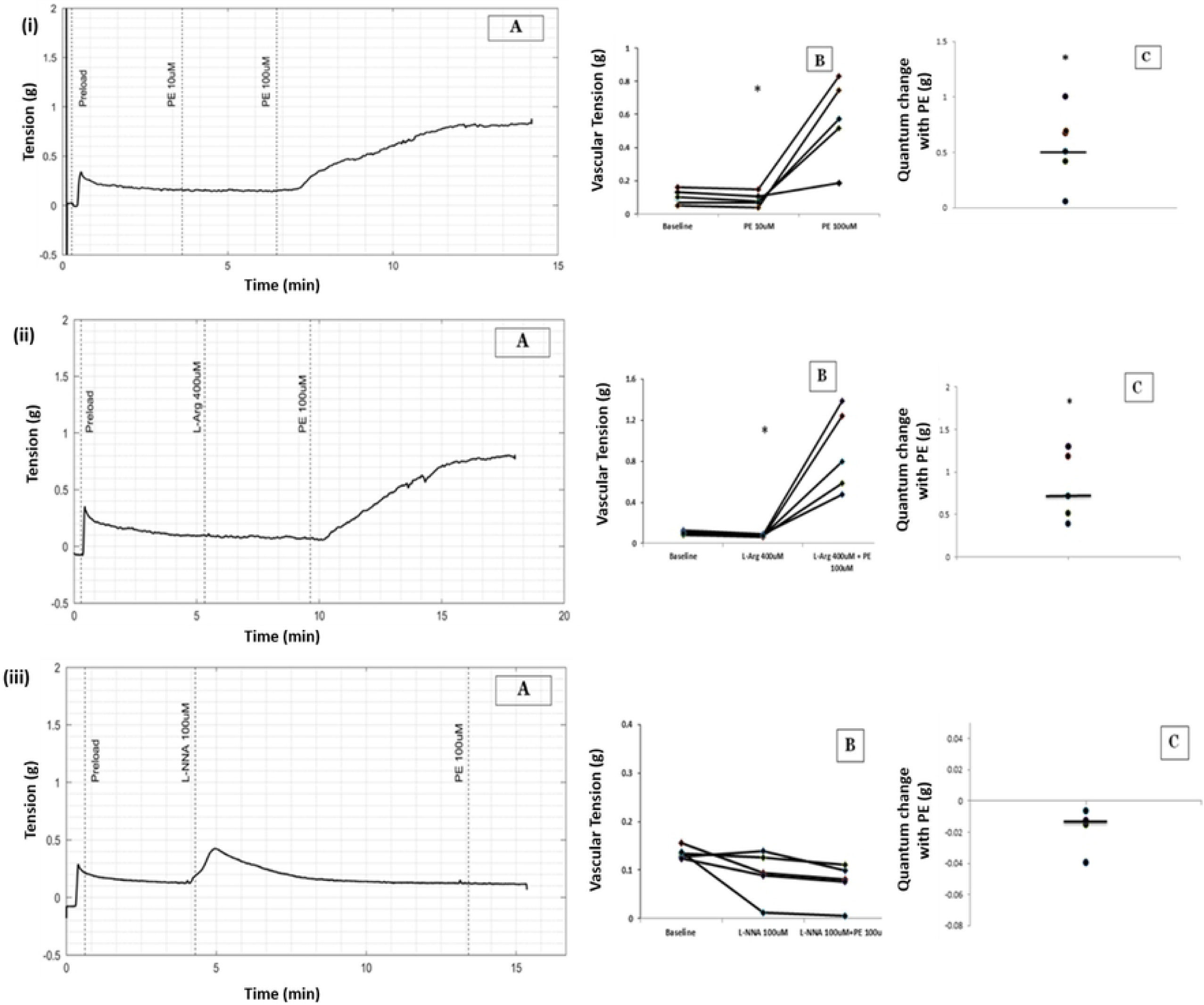
(A) Raw tracings showing changes in vascular tension in longitudinal strips of aorta (B) Scatter plots of results from all five experiments demonstrating increase in vascular tension with PE. (C) Quantum of change with PE from baseline (* p< 0.05). (i) PE 100uM (ii) PE 100uM in high NO environment caused by L-Arginine (iii) PE 100uM in presence of SNP

High NO environment created by addition of L-Arginine 100Um did not cause change in tension of the longitudinal strip of aorta. However subsequent addition of PE 100uM increased tension from 0.22g to 0.32g. (Median, n=5, P value = 0.043, WSR), Fig.3, (ii).

In another set of experiments, in the presence of SNP, a NO donor, PE 100uM increased tension of longitudinal smooth muscle from 0.32g to 0.39g. (Median, n=5, P value = 0.043 with WSR test when tension before and after PE in presence of SNP were compared), Fig.3, (iii).

There was no significant difference when the percentage change in vascular tension due to PE with and without L-Arginine were compared (P value= 0.076 with Mann-Whitney U test, MWU) and also when percentage changes in vascular tension due to PE with and without SNP were compared (P value=0.251 with MWU) showing that PE produced vasoconstriction in aortic longitudinal muscle was not affected by high NO conditions.

### PE induces contraction of circular smooth muscles of small artery under normal NO and high NO environments as expected

10µM concentration of PE did not alter the tension in small-artery preparation, but at 100 µM concentration, PE increased the tension of circular smooth muscle of small artery from 0.09g to 0.54g. (Median, n=5, P value = 0.028 with Wilcoxon signed rank test when tension before and after PE was compared, Fig.4, (i).

**Fig.4:**
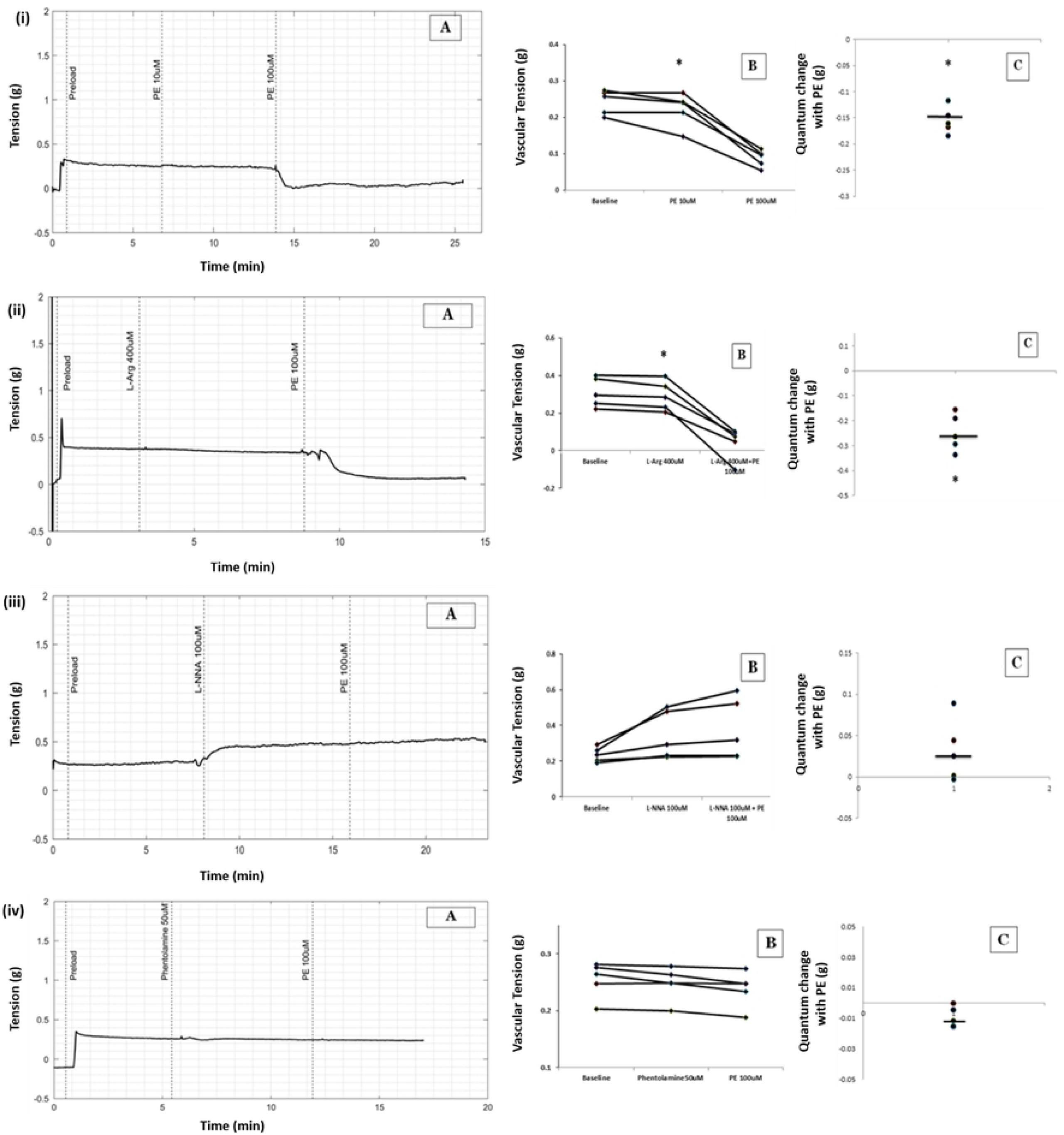
(A) Raw tracing showing changes in vascular tension with PE 100µM in transverse cylinder preparation of small artery. (B) Scatter plots of results from all five experiments demonstrating change in vascular tension with PE. (C) Quantum of change with PE from baseline (* p< 0.05), (i) PE 100uM (ii) PE in the presence of L-Arginine (iii) PE in the presence of NO synthase blocker L-NNA

No change in basal tension was observed when high NO environment was created by adding L-Arginine 100uM. Subsequent addition of PE 100uM increased the tension of circular strip of small artery from 0.07g to 0.79g (Median, n=5, P value = 0.043 with WSR test, Fig.4, (ii).

### L-NNA inhibited PE induced contraction of circular smooth muscle in small artery

Addition of L-NNA a competitive inhibitor of Nitric oxide synthase, caused a transient increase in tension which returned to baseline. Subsequent addition of PE 100µM failed to produce contraction of circular smooth muscle. The vessel tension before and after addition of PE in the presence of L-NNA were 0.09 and 0.08g (median, n=5, P value = 0.08 with WSR test), Fig.4, (iii).

There was a statistically significant difference in percentage of change in tension among the experimental group of PE alone and PE in presence of L-NNA. (Median, n=5, P value = 0.008, with MWU test).

### PE induces relaxation of longitudinal smooth muscle of small artery both under normal and high NO environments

PE 100uM reduced the tension of longitudinal smooth muscle from 0.25g to 0.09g in small arteries. (Median, n=5, P value = 0.043, WSR), Fig.5, (i)).

**Fig.5:**
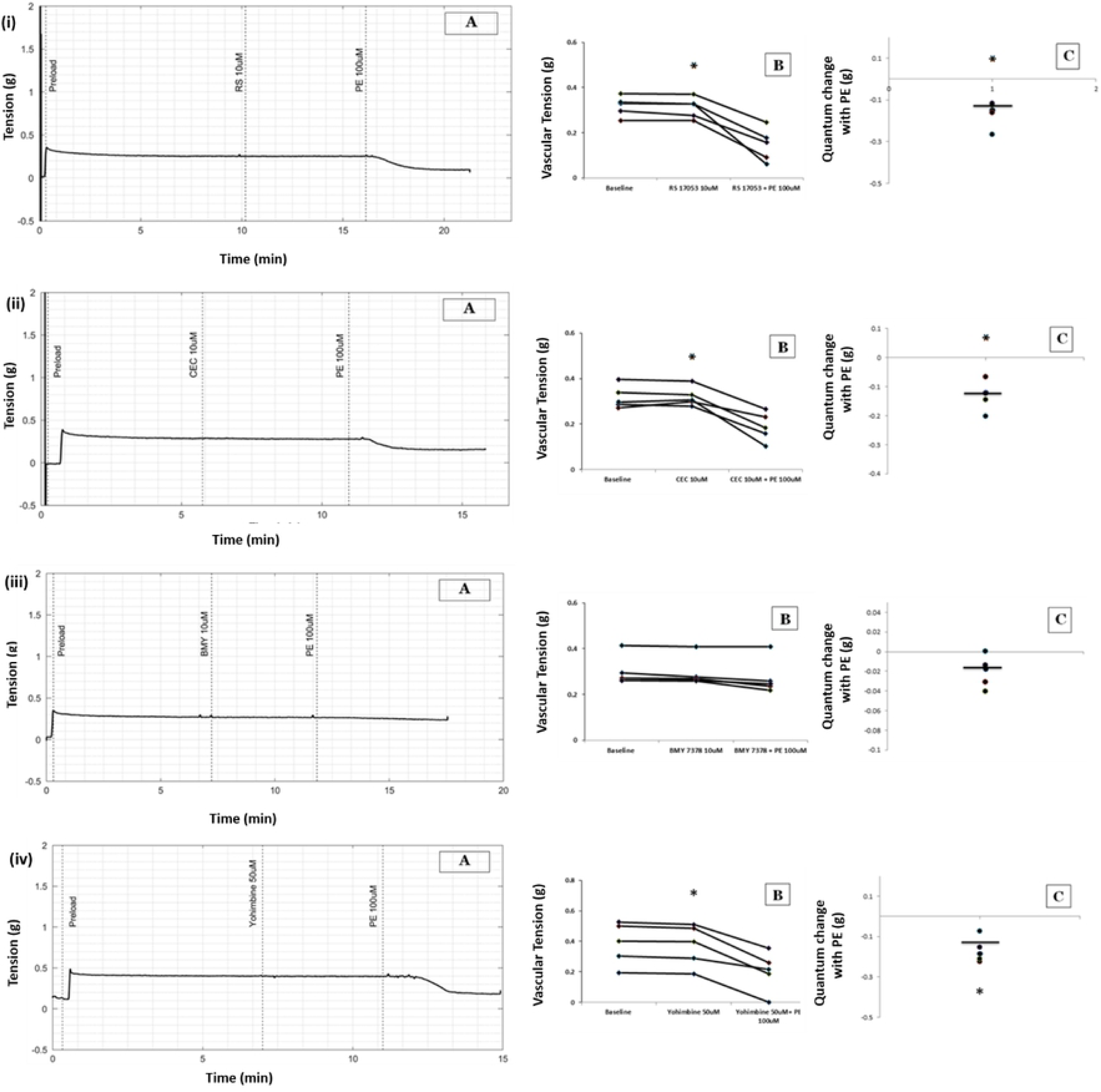
(A) Representative raw tracing showing changes in vascular tension with PE in longitudinal strip of small artery. (B) Scatter plots of results from all five experiments demonstrating changes in vascular tension with PE. (C) Quantum of change with PE from baseline (* p< 0.05), (i) PE 100uM (ii) PE in the presence of L-Arginine (iii) PE in the presence of NO synthase blocker L-NNA (iv) PE in presence of an alpha adrenoceptor blocker, Phentolamine.

While L-Arginine 100uM didn’t reduce the basal tension by itself, subsequent addition of PE 100uM reduced the tension from 0.28 to 0.07g (Median, n=5, P value = 0.043, WSR), Fig.5, (ii).

There was no significant difference in percentage change in tension between the groups: PE under normal NO environment and PE in presence of L-Arginine (P value = 0.08, MWU test).

### PE induced relaxation of longitudinal smooth muscle in small artery was prevented by L-NNA

L-NNA itself increased the basal tension of the vessel. PE 100µM failed to cause relaxation of longitudinal smooth muscle in the presence of L-NNA. The vessel tension before and after addition of PE in the presence of L-NNA were 0.29g and 0.31g. (Median, n=5, P value = 0.13 with WSR test), Fig.5, (iii).

A significant difference in percentage change in tension was observed when the control group (PE alone) and L-NNA/PE group were compared (P value < 0.05, MWU test).

### PE induced relaxation of longitudinal smooth muscle is alpha adrenoceptor mediated

PE failed to produce relaxation of longitudinal smooth muscle in the presence of phentolamine 50uM, an alpha adrenoceptor blocker. The vascular tension before and after the addition of PE (in the presence of phentolamine 50uM) were 0.24g and 0.24g respectively (median, n=5, P value > 0.05, WSR test). Fig.5, (iv).

A significant difference in percentage change of vascular tension was observed when the control group (PE alone) and PE in the presence of phentolamine 50uM was compared (P value < 0.05, MWU test).

### PE induced relaxation of longitudinal smooth muscle is mediated through alpha-1D receptor subtype

Delineation of specific subtype of alpha-1 adrenergic receptor responsible for PE induced vasorelaxation was done by addition of specific subtype receptor blocker. Alpha 1A receptor blocker, RS17053 (10uM) and alpha 1B receptor blocker, CEC (10uM) were not able to prevent the PE induced reduction in tension in longitudinal strip. However, an alpha 1D receptor blocker, BMY 7378 (10uM) prevented PE induced reduction in tension in longitudinal strip.

The vascular tension before addition of PE (in the presence of 10uM RS 17053) was 0.32g and after addition of PE was 0.24g (median, n =5, p value < 0.05 with WSR test), Fig.6, (i). A statistically significant difference was not observed (P value = 0.347, MWU test) when percentage change of vascular tension of the control group (PE alone) and percentage change of vascular tension of PE in the presence of RS 17053 10uM was compared.

**Fig.6:**
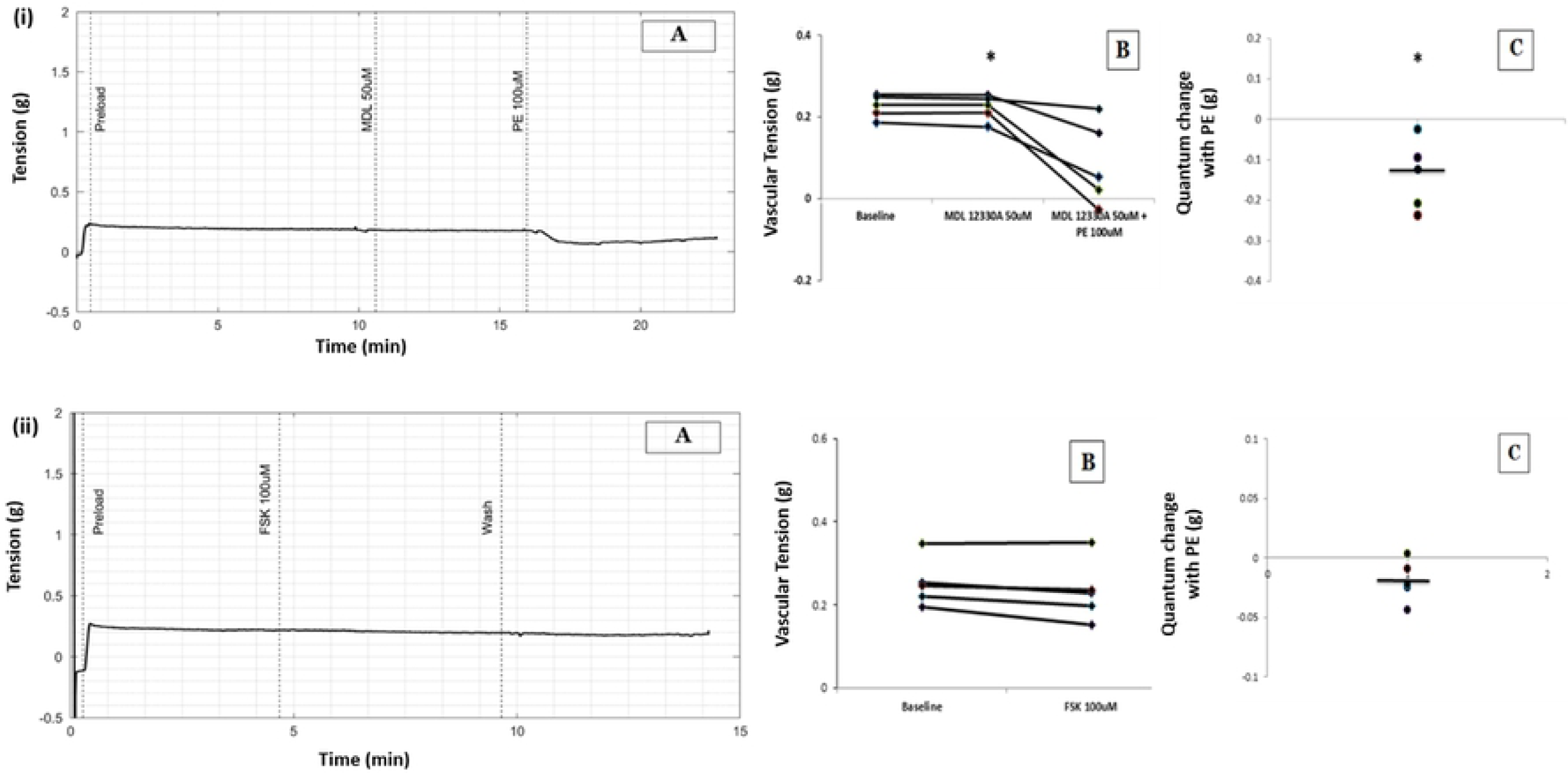
(A) Raw tracing showing changes in tension with PE in the presence of specific blockers in longitudinal strip preparation of small artery. (B) Scatter plots of results from all five experiments demonstrating PE induced changes in vascular tension in presence of specific blockers. (C) Quantum of change PE from baseline, (i) PE in the presence of α1A blocker (ii) PE in the presence of α1B blocker (iii) PE in the presence of α1D blocker (iv) PE in the presence of α2 blocker.

Tension in the longitudinal strip prior to the addition of PE (in the presence of CEC 10uM) was 0.30g and after addition of PE was 0.18g (median, n =5, P value < 0.05 with WSR test), Tension in the longitudinal strip prior to the addition of PE (in the presence of CEC 10uM) was 0.30g and after addition of PE was 0.18g (median, n =5, P value < 0.05 with WSR test), Fig.6, (ii).

There was a 37% reduction of baseline tension by PE in the control group (in the absence of CEC 10uM), (Median, n=5) whereas the reduction of baseline tension in the presence of CEC 10uM was about 56% (Median, n=5). However, for this sample size there was no significant difference when percentage change in tension between the control group and CEC/PE group was compared (P value = 0.056 with MWU test).

The tension in longitudinal strip before addition of PE (in the presence of BMY 7378 10uM) was 0.26g and after addition of PE was 0.24g (median, n=5, P value = 0.08 with WSR test), Fig.6, (iii).

There was a significant difference between the control group and BMY 7378/PE group when the percentage change in tension was compared (P value < 0.05 with MWU test).

### PE induced relaxation of longitudinal smooth muscle is not mediated through alpha-2 receptor

To identify if PE induced vasorelaxation in longitudinal strip of small artery is mediated through alpha-2 adrenergic receptor, yohimbine 50uM, an alpha-2 antagonist was used. Tension in the longitudinal strip prior to the addition of PE (in the presence of Yohimbine 50uM) was 0.39g and after addition of PE was 0.21g (median, n =5, P value < 0.05 with WSR test), Fig.6, (iv).

A significant difference in percentage change of vascular tension was not observed when the control group (PE alone) and PE in the presence of yohimbine 50uM was compared (P value = 0.117, MWU test)

### PE induced relaxation of longitudinal smooth muscle is not mediated through cAMP

To delineate the role of cAMP in PE induced vasorelaxation in longitudinal strips of small artery, MDL 12330A, an adenylyl cyclase inhibitor was used. Tension in the longitudinal strip prior to the addition of PE (in the presence of MDL 50uM) was 0.22g and after addition of PE was 0.05g (median, n =5, P value < 0.05 with WSR test), Fig.7, (i).

**Fig.7:**
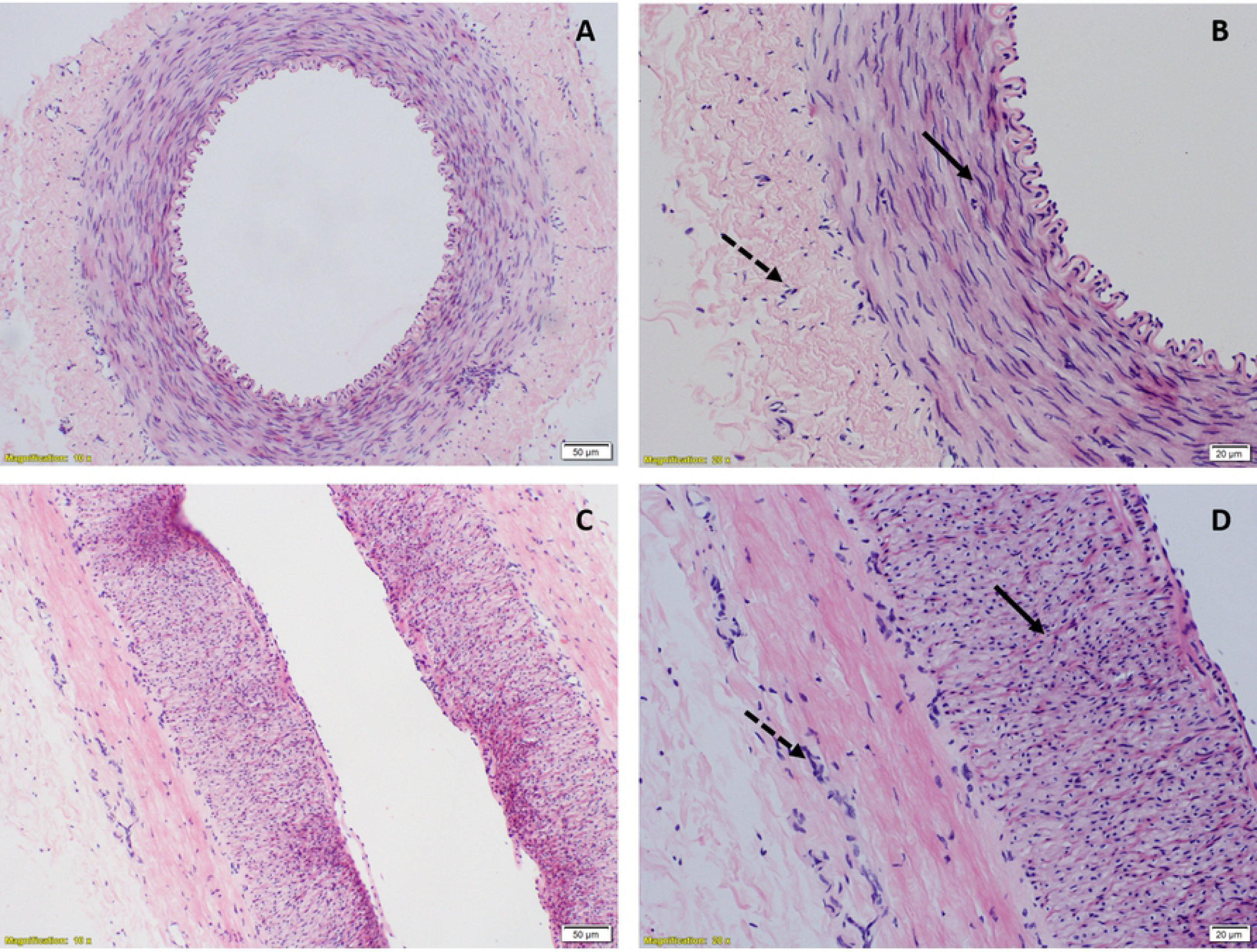
(A) Raw tracing showing changes in tension in longitudinal strip preparation of small artery. (B) Scatter plots showing reduction in vascular tension. (C) Quantum of change with (i) PE in presence of Adenylyl cyclase blocker (ii) Forskolin.

There was no significant difference between the control group (PE alone) and MDL 12330A 50uM group when the percentage change in tension was compared (P value = 1.00, with MWU test).

Forskolin (FSK), an adenylyl cyclase stimulator failed to bring about reduction in tension in longitudinal strips of small artery by itself. Tension in the longitudinal strip prior to the addition of FSK was 0.24g and after addition of FSK was 0.23g (median, n =5, P value = 0.08, with WSR test), Fig.7, (ii).

### Histological Examination

To look for anatomical evidence for presence of longitudinal smooth muscle in small artery, histological examination was done. Light microscopic images of transverse cylinder preparation showed elliptical nucleus of circular smooth muscle predominantly while a few nuclei were seen circular representing a different arrangement of muscle fibres (Fig.8A, 8B). In light microscopic images of longitudinal strip, nuclei of circular smooth muscle were seen circular and a few were seen elliptical representing the longitudinal smooth muscle fibres (Fig.8C, 8D). Similar picture is seen in aorta in which the existence of longitudinal smooth muscle has already been demonstrated (Fig.9A, 9B, 9C & 9D).

**Fig.8:**
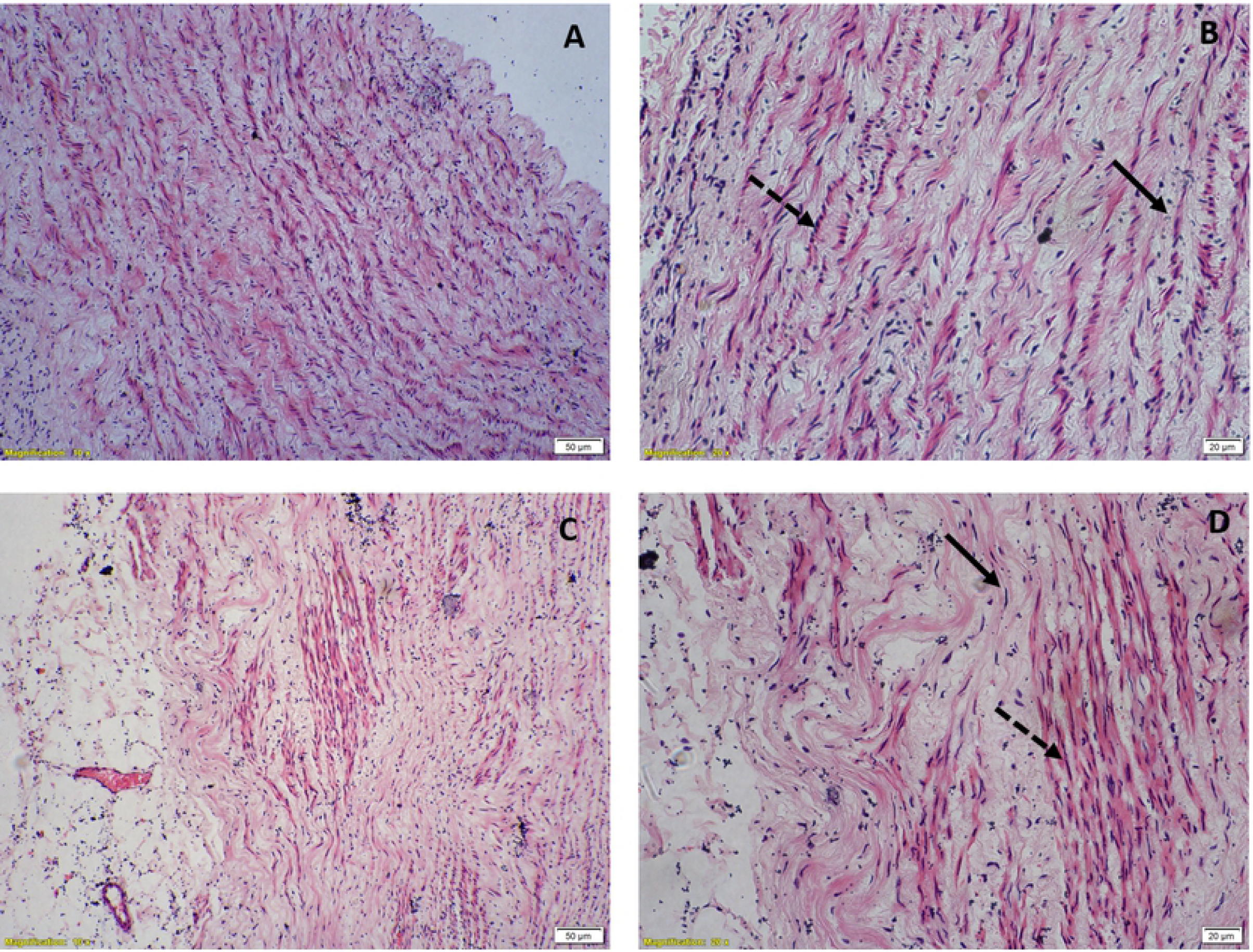
Hematoxylin & eosin stained light microscopic view of small artery. (A) & (B) Transverse cylinder preparation showing elliptical nuclei of circular smooth muscle fibres (solid arrow) and circular nuclei of longitudinal muscle fibres (dashed arrow) under 10X & 20X magnification respectively, (C) & (D) Longitudinal strip preparation showing elliptical nuclei of longitudinal smooth muscle fibres and circular nuclei of circular smooth muscle fibres under 10X & 20X magnification.

**Fig.9:**
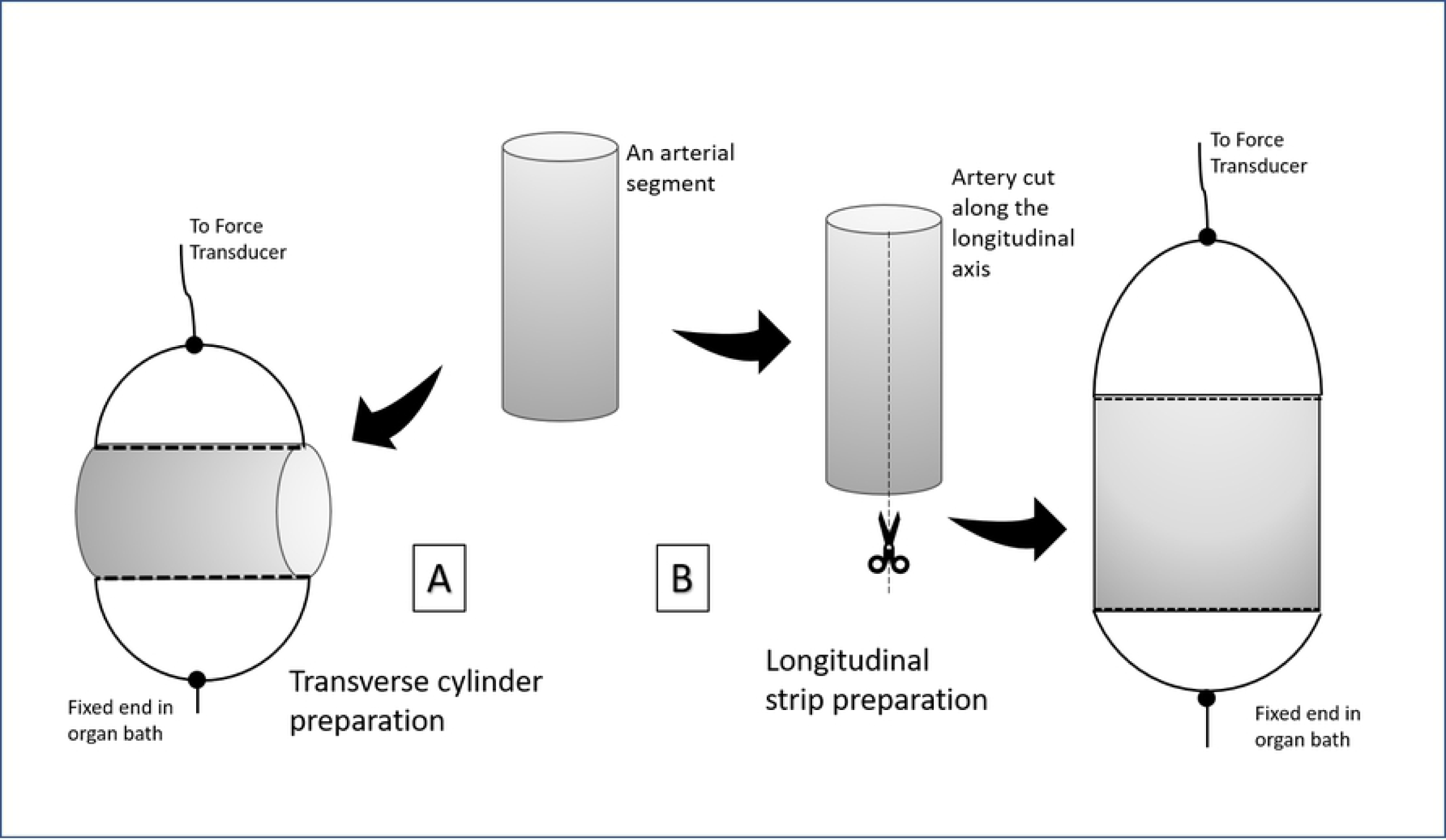
Hematoxylin & eosin stained light microscopic view of aorta, (A) & (B) Circular strip preparation showing elliptical nuclei of circular smooth muscle fibres (solid arrow) and round nuclei of longitudinal muscle fibres (dashed arrow) under 10X & 20X magnification respectively, (c) & (D) Longitudinal strip preparation showing elliptical nuclei of longitudinal smooth muscle fibres (dashed arrow) and round nuclei of circular smooth muscle fibres (solid arrow) under 10X & 20X magnification respectively.

## Discussion

Sympathetic stimulation is known to cause vasoconstriction through alpha adrenergic receptors (6) (7). Adrenergic receptors are broadly classified as alpha adrenoceptors and beta adrenoceptors (8) (9). Alpha adrenoceptors are further classified as alpha 1 and alpha 2 adrenergic receptors (6). Alpha 1 adrenoceptors in turn fall into 3 subtypes namely alpha 1A, alpha1B and alpha 1D. Subtypes of alpha 2 adrenoceptors include alpha 2A, alpha 2B and alpha 2C (6). Beta adrenoceptor subtypes are beta 1, beta 2 and beta 3 (10). All adrenergic receptors are G-protein coupled receptors (9). Alpha and beta 2 adrenoceptors are located on vascular smooth muscle. (11).

Vasoconstriction by alpha adrenergic stimulation occurs in two ways – either stimulation of myosin light chain kinase (MLCK) by calcium/Calmodulin complex (12) or inhibition of myosin light chain phosphatase by (MLCP) by phosphorylated CPI-17 (13) (14)

It has been shown earlier that PE could cause alpha-adrenoceptor mediated vasorelaxation in spiral strips of small arteries in a high nitric oxide (NO) environment (15). However, it was realized that whether a spiral strip will relax or constrict depended not on NO, but in the manner in which it was prepared. There was PE-induced relaxation with strips prepared with fewer spirals (in hindsight this would include more longitudinal component) and PE induced contraction even in a high NO environment with strips prepared in a manner to include more concentric muscle. To clarify this paradox this study focuses on the effect of PE on two different preparations - transverse cylindrical arterial strip (circular smooth muscle) and longitudinal strip (longitudinal muscle) respectively. Both these types of preparations were made with small arteries and aorta. It is shown here that there is a component of longitudinal muscle in small arteries which relaxes on alpha adrenergic stimulation. However, circular strips of small arteries demonstrated the well-known vasoconstrictor response with alpha adrenergic stimulation (with PE). The relaxant response of longitudinal smooth muscle to PE that was observed in small arteries did not occur in the aorta. Both circular and longitudinal preparations of aorta showed a vasoconstrictor response with PE.

Phenylephrine, a selective alpha agonist (16) is a known vasoconstrictor, the effects of which are mediated via alpha 1 and alpha 2 adrenergic receptors. It has been used as a vasopressor agent in management of hypotensive states of various etiologies. Phenylephrine acts on G_q_ coupled alpha 1 receptor which itself is a GTPase. The alpha subunit of G_q_ gets cleaved and activates phospholipase-C on the cell membrane(17). Phospholipase-C causes breakdown of phosphatidylinositol-4,5 bisphosphate (PIP2) on cell membrane to inositol 1,4,5-triphosphate (IP_3_) and 1,2 -diacylglycerol (DAG). IP_3_ then enters the cytoplasm, acts on IP3 receptors on sarcoplasmic reticulum and causes release of calcium. The released calcium then binds to calmodulin, a regulatory protein forming calcium/calmodulin complex (18). Calcium/Calmodulin complex causes stimulation of MLCK which causes phosphorylation of myosin light chain thereby causing contraction (12). Calcium released from sarcoplasmic reticulum combines with DAG and stimulates protein kinase C (PKC). PKC causes phosphorylation of a smooth muscle phosphoprotein, CPI-17. CPI-17 inhibits MLCP, which prevents dephosphorylation of myosin light chain thereby supporting contraction (13) (14). Activation of alpha 2 adrenoceptor stimulates G_i_ type of G-protein, which in turn inhibits adenylyl cyclase and decreases cAMP, again supporting contraction (18).

Raj et al., (15) have demonstrated in spiral strip preparation of goat arteries, scenarios in which PE could cause vasorelaxation from basal tone, one being high NO environment. It was shown that PE/NO induced vasorelaxation is cGMP independent and requires alpha adrenergic activation. However, further experiments showed that contractile or relaxant response depends on the manner of spiral strip preparation rather than on the level of NO.

The results show that PE produces only contraction of circular smooth muscle both under normal NO and high NO environment as known earlier. However, the effect of PE on longitudinal component of smooth muscle is only relaxation both under normal NO and high NO environment. L-Arginine is reported to produce relaxant response by increasing NO levels (19). However, we observed that L-Arginine by itself did not induce relaxation in both small artery and aorta. It is not as if the preparation were not capable of relaxation because in the case of longitudinal strip of small artery at least PE could induce vasorelaxation. The earlier observations of L-Arginine induced relaxation were all made in preconstricted strips (20).

It was also found that such PE induced vasorelaxation in longitudinal smooth muscle is alpha adrenergic dependent as phentolamine, an alpha adrenoceptor antagonist prevented such response. In order to delineate the receptor subtype involved in PE induced vasorelaxation, four alpha adrenoceptor α_1A_, α_1B_, α_1D_ and α_2_ subtype blockers were used. Among them only α_1D_ blocker, BMY 7378 was successful in preventing PE induced vasorelaxation. BMY can block alpha-2C adrenoceptors when used at a concentration above 0.3 μM (21). Though the concentration that we have used is 10 μM, its effect of blocking vasorelaxation in longitudinal strips is unlikely to be due to alpha 2C blockade, as Yohimbine, which is a non-selective antagonist or alpha-2 subtypes did not prevent the vasorelaxation induced by PE. Hence, prevention of PE - induced vasorelaxation by BMY 10 μM is through alpha-1D receptor.

The role of second messenger, cAMP in PE induced vasorelaxation was also studied. To decipher this forskolin, adenylyl cyclase activator and MDL 12330A, adenylyl cyclase inhibitor were used. Forskolin by itself failed to produce vasorelaxation in longitudinal strip and MDL 12330A also failed to prevent PE induced vasorelaxation showing that PE induced vasorelaxation in longitudinal strip is not mediated through cAMP.

It is also shown that L-NNA, a competitive inhibitor of NO synthase inhibited both PE induced contraction and relaxation in small artery. The probability of L-NNA being a blocker of alpha-adrenergic receptors too rather than being a NO synthase blocker has to be evaluated further. Also, L-NNA caused a transient increase in tension in transverse cylinder preparation whereas it produced a sustained increase in tension in longitudinal preparation of small artery. This differential behaviour of L-NNA also needs further evaluation.

Further, anatomical evidence for the existence of longitudinal smooth muscle in small arteries has been demonstrated through histology.

Presence of longitudinal smooth muscle in aorta is well-known. However, the effect of PE on both circular and longitudinal smooth muscle of aorta is only vasoconstriction, unlike in small arteries.

This is the first time within our knowledge, the existence of longitudinal smooth muscle in small arteries is being reported. It is also shown here that the longitudinal smooth muscle of small artery relaxes in response to α_1D_ adrenergic stimulation and such relaxant response is independent of cAMP. However, the physiological significance for the existence of such longitudinal component of smooth muscle in small arteries and their relaxant response to alpha adrenergic stimulation needs to be ascertained.

